# Quantitative contribution of the spacer length in the supercoiling-sensitivity of bacterial promoters

**DOI:** 10.1101/2022.02.25.481925

**Authors:** Raphaël Forquet, William Nasser, Sylvie Reverchon, Sam Meyer

## Abstract

DNA supercoiling acts as a global transcriptional regulator in bacteria, but the promoter sequence or structural determinants controlling its effect remain unclear. It was previously proposed to modulate the torsional angle between the -10 and -35 hexamers, and thereby regulate the formation of the closed-complex depending on the length of the “spacer” between them. Here, we develop a thermodynamic model of this notion based on DNA elasticity, providing quantitative and parameter-free predictions on the relative activation of promoters containing a short vs long spacer when the DNA supercoiling level is varied. The model is tested through an analysis of *in vitro* and *in vivo* expression assays of mutant promoters with variable spacer lengths, confirming its accuracy for spacers ranging from 15 to 19 nucleotides, except those of 16 nucleotides where other regulatory mechanisms likely overcome the effect of this specific step. An analysis at the whole-genome scale in *E. coli* then demonstrates a significant effect of the spacer length on the genomic expression after transient or inheritable superhelical variations, validating the model’s predictions. Altogether, this study shows that the torsional constraints associated to promoter binding by RNA Polymerase underpin a basal and global regulatory mechanism independent of specific transcription factors.

## Introduction

DNA supercoiling (SC), the level of over- or underwinding of the double-helix, is a fundamental property of DNA. In bacterial cells, the chromosome is maintained at a negative SC level (in an underwound state), by a finely controlled balance between the DNA relaxation activity of topoisomerase I (and IV), and the introduction of negative supercoils by the DNA gyrase [1, 2]. This level is affected by a variety of factors, including growth phase and environmental conditions [3]. In addition to playing a key role in the physical organisation of the chromosome [4], SC was soon discovered to affect the expression of many promoters [5, 6]. More recently, through the advent of transcriptomic technologies allowing genome-wide expression profiling, it was shown to act as a global transcriptional regulator in many bacteria, based on studies employing gyrase inhibitors inducing a global DNA relaxation and, in turn, a global and complex response of genes [3]. Fast changes in SC levels may thus play an important and global role in the transcriptional response of bacteria to environmental changes [1, 2].

At the mechanistic level, SC affects the transcription process at multiple steps, both indirectly through regulatory proteins, and directly by modulating the interaction of RNA Polymerase (RNAP) with DNA [3]. This includes the formation of the open-complex during transcription initiation, which is strongly facilitated by negative SC [7, 8] and plays a key role in the SC response of promoters [9]. But other steps of transcription also contribute to this response, including closed-complex formation [10], promoter escape [11], transcription elongation and termination [12]. Because of this complexity, regulatory models able to predict the response of a given promoter to SC variations are mostly lacking, at the quantitative and even the qualitative levels [3, 13], in contrast to those involving regulatory proteins [14]. Considering the widespread relevance of this mechanism in bacterial gene expression, obtaining such quantitative models is an important objective, both for fundamental understanding of the process and for applications, e.g., in synthetic biology [15]. The objective of this paper is to develop such a model, focusing on the specific step of closed-complex formation during transcription initiation.

The latter step involves the binding of RNAP to gene promoters, through specific recognition of the -35 and -10 elements by 2.4 and 4.2 regions of the *σ* factor respectively [16]. The efficiency of RNAP binding on the promoter depends primarily on the proximity of -35 and -10 sequences to their respective consensus, but also on the spacer element between them. The latter exhibits little or no contact with the transcription machinery, except for promoters with an extended -10 element which interacts with the 3.0 region of *σ* factors [16]. The spacer exhibits a variable length, ranging usually from 15 to 19 nucleotides (nt) for *σ*70-dependent promoters [16, 17]. For some promoters, the spacer is ill-defined because the -35 sequence is weak, which is then compensated by an extended -10 element [18] or by the assistance of regulatory proteins such as CRP [19]. The maximal core promoter activity is reached with spacers of 17 nt, while the addition or subtraction of nucleotides from this optimal length reduces their expression by several-fold [20, 21]. While the spacer sequence exhibits no specific requirement, point mutations [22, 23] or modifications of its AT richness [24, 25] also affect promoter expression [15, 26], presumably by altering its 3D conformation [27, 28, 29].

In addition to altering promoter strength, the spacer length has been shown to strongly modulate their SC response [20, 21]. Accordingly, most promoters of stable RNAs in *E. coli*, which are strongly repressed by DNA relaxation, were found to contain spacers of unusual length (16 nt) [10]. A qualitative model of this regulation mode was proposed around 30 years ago, based on a simple geometric effect [10]. Because of the helical nature of DNA, variable spacer lengths are associated to different relative orientations of the -35/-10 binding sites, which might modulate RNAP binding. In turn, the presence of torsional stress in the spacer might rotate the -35/-10 binding sites toward a (un)favourable orientation and thus regulate RNAP activity (Fig. 1A). This qualitative notion was supported by a review of observations from a collection of individual promoters [10, 30, 15], and was also suggested as the mechanism of activation of several TFs [31]. According to this mechanism, promoters containing suboptimal spacer lengths are not simply under-expressed, but might rather be selectively down- or up-regulated by the cell according to their length, depending on the global SC level.

**Figure 1:**
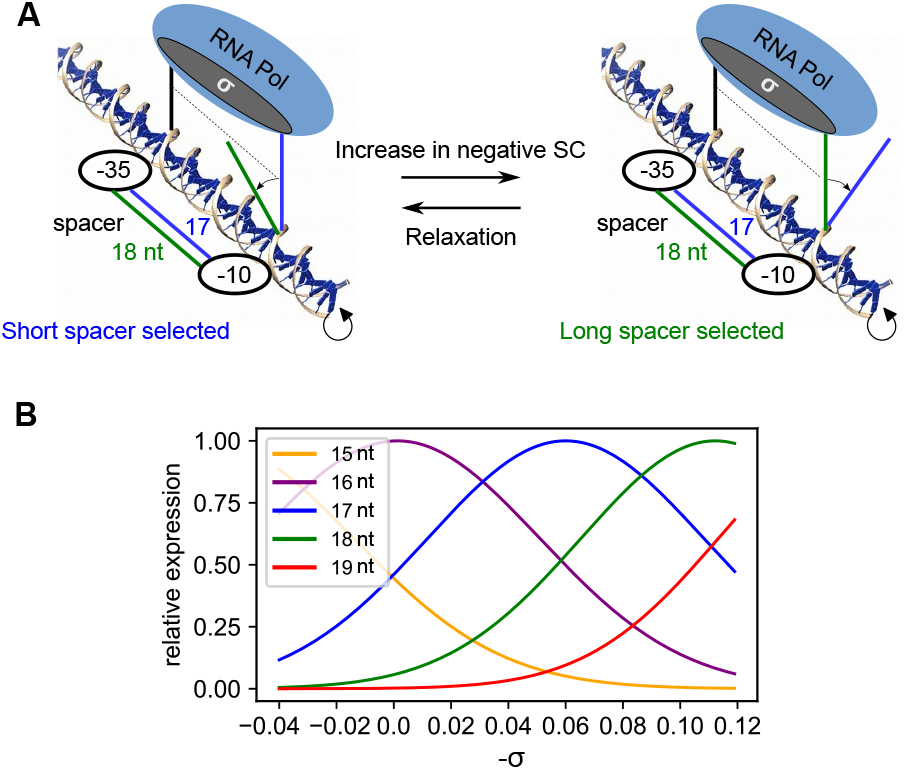
Quantitative modelling of the coupling between RNAP binding sites orientation, spacer length and DNA supercoiling. **(A)** Schematic depiction of -35/-10 alignment depending on spacer length and superhelical density. With long spacers (green), RNAP binding sites are out-of-phase at moderate SC levels, and become optimally aligned when DNA is highly negatively supercoiled, and conversely for shorter spacers (17-nt spacer in blue). **(B)** Relative expression levels predicted by the regulatory model for 15 to 19-nt spacers, depending on *σ*. Short spacers are more expressed at relaxed levels, while long spacers are favoured at highly negative levels. The only parameter *θ*_*P*_, representing the optimal angle for RNAP binding, was chosen such that 17-nt spacers are maximally expressed at *σ* = *−*0.06 (see text).

In this study, we propose to (1) translate this qualitative notion into a quantitative regulatory model, (2) develop a rigorous validation of its predictions, based on *in vitro* and *in vivo* analyses of mutant promoters containing variable spacer lengths and (3) demonstrate the genome-scale relevance of this mechanism by a statistical analysis of high-throughput expression data, which were not available when the previous qualitative models were developed but are particularly suited to this global regulation mode. These results consistently show that the variability of spacer lengths underpins a global selectivity of promoter activity depending on the cellular SC level, varying either transiently due to topoisomerase inhibitors, or inheritably in the longest-running evolution experiment. They confirm and provide a quantitative framework to the proposed basal regulatory mechanism, which might play a widespread role in the prokaryotic kingdom.

## Materials and Methods

### Thermodynamic model of supercoiling-dependent transcription

We assume that the formation of the closed-complex is limited by an intermediate state where the spacer DNA must be (un)twisted to a favourable relative orientation of the -10 and -35 sites, allowing RNAP binding (Fig. 1A). This deformation is treated in the elastic approximation, and the associated orientational free energy depends on the spacer length *n* and average superhelical level *σ*:

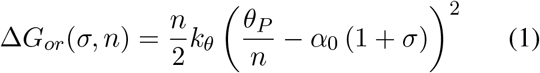

where *k*_*θ*_ = 71.4 *k*_*B*_*T*.*rad*^−2^ is the DNA sequence-averaged twist stifness [32, 33], *k*_*B*_*T* is the Boltzmann factor, *α*_0_ = 34° is the average twist angle between adjacent nucleotides [34, 35], and *θ*_*P*_ is a global parameter representing the optimal twist angle between - 35 and -10 sites for RNAP binding. To illustrate the model predictions (Fig. 1) and subsequent calculations, we assumed that 17-nt spacers achieve this optimal angle at the standard superhelical level *σ* = *−*0.06, i.e. *θ*_*P*_ = 17 *× α*_0_ *×* (1 *−* 0.06) = 543°, but most key predictions are independent of its value (see upcoming paragraphs).

In this calculation, we assume that the total superhelicity *σ* can be utilised by RNAP for the twist deformation of spacer DNA (although most of it is stored as writhe at equilibrium). This unidimensional approximation of DNA is frequently used in analyses of the regulatory effect of SC [36, 10], and is relevant here in particular because the spacer DNA is too short to significantly writhe (before RNAP binding). Since RNAP is assumed to impose a fixed angle *θ*_*P*_, its flexibility is neglected in the calculation.

The total free energy associated to the transcription process is assumed to contain three contributions related to the effects of SC, of the spacer element, or of both simultaneously:

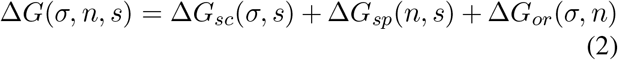

where

- Δ*G*_*or*_(*σ, n*) is the orientational deformation energy (Eq. 1), and introduces a *coupled* dependence on *σ* and *n*.
- Δ*G*_*sc*_(*σ, s*) depends on the promoter sequence *s*, and represents all other mechanisms of regulation by SC (e.g., promoter opening, 3D deformations, structural transitions, etc.), which are assumed to be independent of spacer length *n*.
- Δ*G*_*sp*_(*n, s*) represents all other mechanisms of modulation of transcriptional activity by the spacer DNA, which depend on its sequence as well as its length (e.g., stretching, 3D conformation, sequence-specific interactions with RNAP within the closed-complex, etc), but are assumed to be independent of *σ*.

Transcription rates are then computed using a standard thermodynamic framework [14]:

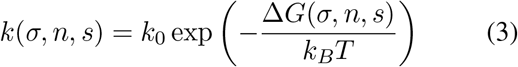

where *k* is the transcription rate, *k*_0_ is the basal rate (which depends, e.g., on the -10/-35 sequence affinities for RNAP), and *k*_*B*_*T* is the Boltzmann factor. To simplify the notations, we use the latter as the energy unit in the following. Based on the equations above, the model does not predict the general SC-dependence of a promoter nor the effect of the spacer on the absolute expression level (since both depend on unpredictable terms in Eq. 2), but it does predict how the SC-sensitivity depends on spacer length by Eq. 1. Since most of the parameters involved in the latter are known from physical measurements, it is possible to derive several quantitative and parameter-free predictions underpinning all analyses of the manuscript, as follows.

### Prediction of relative *in vitro* expression levels depending on spacer length

From *in vitro* expression data of mutant promoters (Fig. 2), we isolate the specific effect of the spacer length *n* by normalising each datapoint by the corresponding value obtained in the reference mutant promoter (*n*_0_ = 17, thereby eliminating any other regulatory effect of SC:

**Figure 2:**
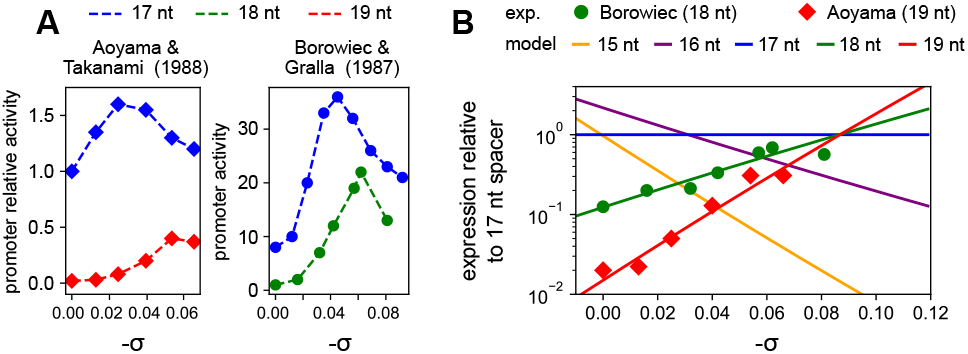
Comparison of *in vitro* transcription assays to model predictions. **(A)** Left panel: relative activities of promoters with 17 and 19-nt spacers (blue and red, respectively) depending on SC level *σ*, data from [46]. Right panel: activity of promoters with 18-nt and 17-nt spacers (green and blue, respectively, data from [45]). **(B)** Transcription model predictions (solid lines) compared to the normalised *in vitro* promoter expression data from [46] (red diamonds) and [45] (green dots). Each datapoint was divided by the corresponding value obtained in the reference mutant promoter with a 17-nt spacer (see text). The slopes of the lines are parameter-free and proportional to *n −* 17 (where *n* is the spacer length), while the intercepts are promoter sequence-specific (see Materials and Methods).

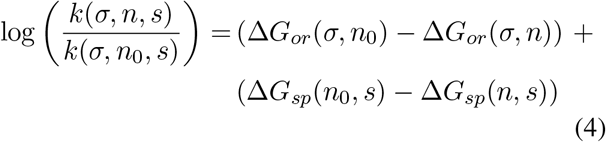

The second term is independent of *σ*, and is thus a constant for a promoter of given sequence and spacer size, hereafter quoted *Q*_*sp*_(*n, s*). Using a linear expansion in Δ*n/n*_0_ = (*n − n*_0_)*/n*_0_ in the first term, the relative expression level of each spacer length simplifies to a linear dependence in *σ* (as visible in Fig. 2B without approximation):

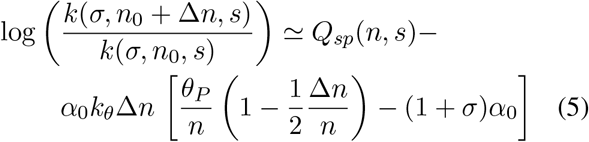

Crucially, the *slope* of each line in Fig. 2B (depending on *σ*) is therefore defined without any adjustable parameter, and is proportional to the torsional stiffness of DNA and to Δ*n* = *n −* 17. The intercept depends on the global parameter *θ*_*P*_, and may also depend on the spacer sequence and length due to *Q*_*sp*_(*n, s*).

### Prediction of *in vivo* expression fold-changes during superhelical variations

All analysed *in vivo* data (from mutant promoters or transcriptomics data) involve relative expression levels (fold-changes) induced by a global superhelical variation *σ*_0_ → *σ*_0_ + Δ*σ* (induced by antibiotics or mutations). The predicted value simplifies to:

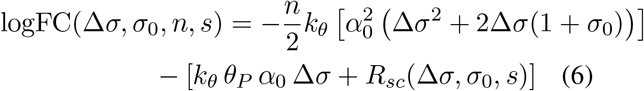

where *R*_*sc*_ = Δ*G*_*sc*_(*σ*_0_ + Δ*σ, s*) *−* Δ*G*_*sc*_(*σ*_0_, *s*) reflects all spacer length-independent regulatory effects of SC on the considered promoter *s*. Crucially, only the first term of the equation depends on the spacer length *n*, and it does not depend on any unknown parameter. Thus, after linear expansion in Δ*σ*, the *relative* effect of the superhelical variation on spacer length mutants of the same promoter *s* is entirely predictable and independent of *σ*_0_ (which is satisfactory since the absolute SC levels are not always known with precision in the analysed *in vivo* data):

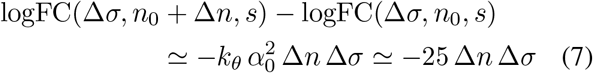

This dependence is shown in Supplementary Fig. S2 for a DNA relaxation Δ*σ* = 0.03, and yields an expression ratio of around 2 between spacers differing by one nucleotide. In transcriptomic analyses, promoters are grouped by their spacer length *n* but differ by their overall sequence *s*, so that the term *R*_*sc*_ remains in the form of strong statistical noise (Fig. 4 and 5), imposing to work with the proportion of activated promoters rather than directly with the fold-change values.

### Measurement of mutant promoters’ responses to opposite supercoiling variations *in vivo*

230 bp sequences upstream of *pheP* start codon were synthesised with mutations in the spacer (GeneCust), and individually cloned into pUCTer-luc plasmids upstream of a luciferase reporter gene (*luc*) (Supplementary Tab. S1). *E. coli* strain MG1655 cells were then transformed with these plasmids using a standard electroporation procedure. The following protocol is described elsewhere [9]. Briefly, *E. coli* cells carrying the plasmids with the different promoters were grown at 37°C in LB medium in a microplate reader (Tecan Spark). The *OD*_600*nm*_ and luminescence were measured every 5 minutes to fol-low bacterial growth and promoter expression, respectively. DNA relaxation was induced by injecting 5 μL of novobiocin (50, 100, 150 and 200 μg/mL final concentrations tested), whereas DNA overtwisting was induced by injecting 2 μL of seconeolitsine (25, 50, 75 and 100 μM final concentration tested) [37]. The responses to such opposite DNA SC variations were then computed by comparing the luminescence values (in triplicates) of the novobiocin or seconeolitsine-shocked strain compared to the same strain injected with water (novobiocin solvent) or DMSO (seconeolitsine solvent), 60’ or 5’ after shock, respectively. The employed firefly luciferase has a lifetime of around 45’ in *E. coli* [38], and buffers the repressive effect of novobiocin. Confidence intervals and p-values were computed using Student’s distributions. Raw datapoints are provided in Supplementary Fig. S3. We previously showed [9] that the presence of the employed plasmids does not affect bacterial growth, and that results are not affected by plasmid copynumber variations (we compute relative expression levels between mutant promoters rather than absolute levels, and consistent observations were obtained in plasmid-borne or in chromosomal luciferase fusions), in agreement with other studies [39, 40]. The employed plasmids are well established as reflecting the average SC level of the chromosome [41], and also specifically in response to novobiocin-induced relaxation [42, 43, 44].

### Parameter fitting in mutation data

In Fig. 1B and 2B, for simplicity, the curves were drawn based on the orientational contribution (Eq. 1) only, with the value *θ*_*P*_ = 543°. In Fig. 2B, the datapoints of [45] fall on the predicted line without adjustment, suggesting *Q*_*sp*_ = 0 for these promoters (see Eq. 5). The data of [46] were fitted with a value *Q*_*sp*_(*n, s*) = 1.1 *k*_*B*_*T* (Eq. 5), corresponding to a factor 3 in expression.

Expression fold-changes measured in microplates with *pheP*-derived promoters were reproduced (Fig. 3), starting from a level *σ* = *−*0.06, with an overtwisting magnitude Δ*σ* = *−*0.02 (seconeolitsine), and a relaxation magnitude Δ*σ* = *−*0.005 (novobiocin). This lower relaxation magnitude probably partly reflects a buffering effect of the reporter system and should thus be considered as an effective value used in the modelling, as further suggested by the lower repressive effect of novobiocin compared to batch cultures [47]. The spacer-independent effect of SC, *R*_*s*_*c* (Eq. 6), was estimated from the repression (novobiocin) and activation (secone-olitsine) level observed with the 17-nt spacer, with values *R*_*sc*_ = 0.4 *k*_*B*_*T* and *R*_*sc*_ = − 0.97 *k*_*B*_*T* corresponding to activation factors of 0.67 and 2.1, respectively (Fig. 3, 17-nt spacers). These presumably reflect the modulation of promoter opening energy by the two opposite superhelical variations [9].

**Figure 3:**
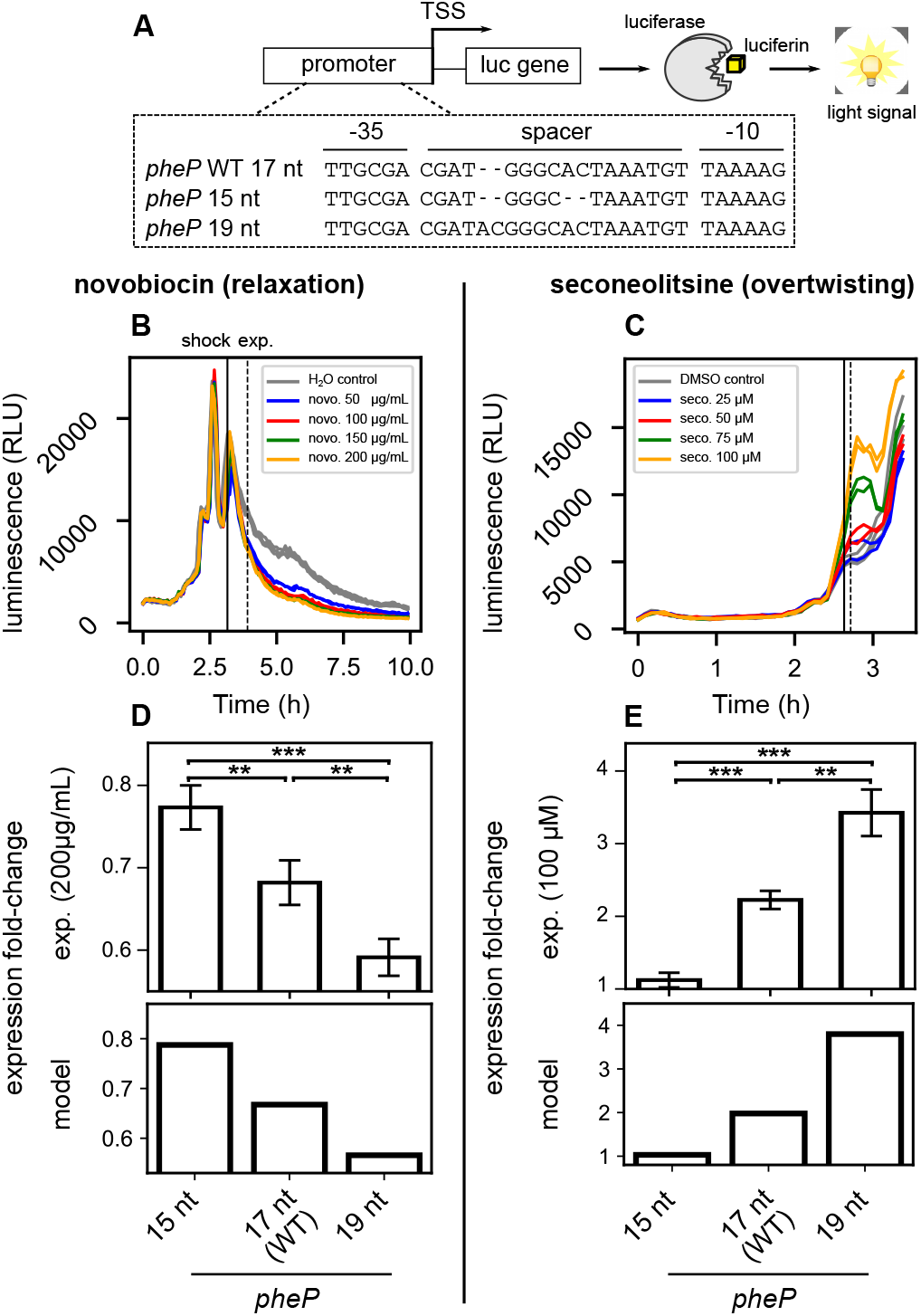
Responses of *pheP*-derived mutant promoters of variable spacer length to opposite SC variations. **(A)** Promoter sequences were synthesised from *pheP* (*E. coli*), with mutated spacers of different lengths. They control the expression of a *luc* gene encoding firefly luciferase, generating luminescence from luciferin substrate. **(B)** Promoter expression monitored in a microplate reader (bacteria carrying plasmids with *pheP* native promoter in LB medium), with a novobiocin shock applied in mid-exponential phase (time point quoted “shock”) at different sublethal concentrations. **(C)** same as (B), with a seconeolitsine shock performed at different sublethal concentrations, which induces a stronger and more transient expression variation. **(D-E)** Expression fold-changes computed 60’ (time point quoted “exp.”) after novobiocin shock (200 μg/mL), or 5’ after seconeolitsine shock (100 μM) from the experiments (upper panels), or predicted by the model (lower panels) assuming SC variations compatible with the observed expression variations levels (see Materials and Methods). All raw datapoints are in Supplementary Fig. S3.

### Genome-wide analyses of spacer responses to supercoiling variations

Transcriptomic responses to DNA relaxation or inheritable SC variations were collected from the literature (Supplementary Tab. S2). The curated map of *E. coli* promoters with associated genes and spacer lengths was retrieved from Ecocyc [48]. Only *σ*70-dependent promoters were retained and classified depending on their spacer length and response to the investigated condition, under standard statistical selection procedures (adjusted *P* -value < 0.05). For transcriptomes of *E. coli* evolved strains, a less stringent *P* -value threshold (0.1) was applied to have enough statistical power for the analysis [49]. For other species, TSS maps were retrieved from the literature [50, 51, 52, 53], and the positions of promoter elements were predicted using bTSSfinder [54]; details are given in Supplementary Information. The relation between promoter activation and spacer length was quantified either by a Student’s t-test between activated and repressed promoters (Fig. 4A, Fig. 5BC, Supplementary Fig. S4) or by linear regression (Fig. 4B). All computations were carried using a homemade Python package. All error bars shown are 95% confidence intervals.

**Figure 4:**
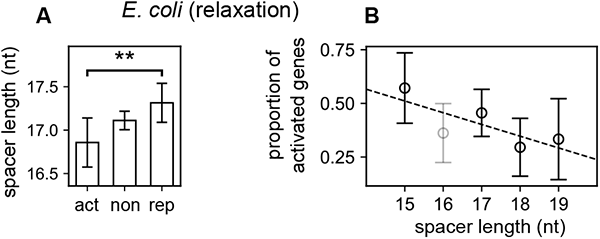
Genome-wide relation between spacer length and promoter selectivity during DNA relaxation in *E. coli*. **(A)** Comparison of the mean spacer lengths of *σ*70-dependent promoters activated (act), not significantly affected (non) or repressed (rep) by norfloxacin-induced DNA relaxation (LZ54 vs LZ41 strains [47]). As expected from our modelling for a DNA relaxation, activated promoters have significantly shorter spacers compared to repressed ones (*P* = 0.007). **(B)** Proportion of activated promoters among those responsive to DNA relaxation, depending on their spacer length (linear regression *P* = 0.07). As observed in *in vitro* data above (Supplementary Fig. S5), 16-nt spacers (in grey) do not follow the model and were excluded from statistical analyses (see Discussion).

**Figure 5:**
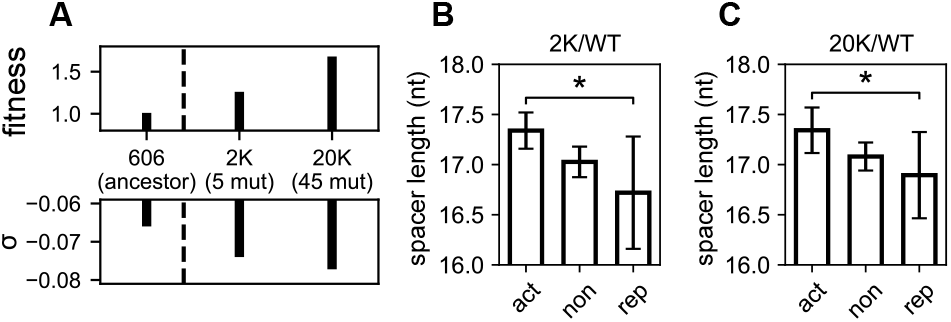
Contribution of the spacer to the global selectivity of promoters by inheritable increase of negative supercoiling in *E. coli*. **(A)** Reproduced from [49]. Strains from the longest-running evolution experiment [58]: 606 (ancestral genetic background), 2K (clone isolated from population at 2,000 generations), 20K (at 20,000 generations). Evolved strains exhibit higher chromosomal SC density *σ* compared to the ancestor, due to the natural acquisition of two point mutations: one in *topA* before 2,000 generations (among the five observed), and one in *fis* before 20,000 generations (among the 45 observed). Those mutations are associated to fitness gains through global expression changes [49]. **(B)** Comparison of mean spacer lengths for promoters activated (act), not significantly affected (non) or repressed (rep) in the 2K evolved strain compared to the ancestor. As expected from our modelling for an increase of negative SC, activated promoters have longer spacers compared to repressed ones (*P* = 0.04). **(C)** Same for the 20K evolved strain compared to the ancestor, where the same difference is observed (*P* = 0.026). The wider confidence intervals for repressed promoters result from their lower number in the investigated conditions.

## Results

### Regulatory model of -35/-10 alignment during closed-complex formation

Following previous works [10, 30], we hypothesised that, for a simultaneous binding of the -35 and -10 hexamers by the RNAP holoenzyme during closed-complex formation, an intermediate state must be achieved where the spacer DNA is (un)twisted to a favourable orientation (Fig. 1A). We translated this notion into a quantitative regulatory model, based on a thermodynamic description of transcription [14] where this contribution can be computed from the torsional energy of spacer DNA, while RNAP is assumed to impose an optimal angle between the two sites. Treating DNA as a homogeneous polymer of known torsional stiffness, this energy depends on the spacer length and its superhelical state before RNAP binding, and the only adjustable parameter is the optimal angle of RNAP (Fig. 1A, detailed hypotheses and equations are given in Materials and Methods).

Since SC affects transcriptional activity at many other steps of the process (promoter opening, escape, elongation, etc.), we assume these other regulatory effects to be independent of the spacer length. Similarly, all other mechanisms by which the spacer length and sequence modulate transcriptional activity (e.g., DNA stretching, 3D deformations, specific interactions with RNAP within the closed complex, etc.) are assumed to be independent of the superhelical state of the promoter preceding RNAP binding. Based on these simplifying assumptions, the modulation of the torsional angle between the -35 and -10 hexamers is the only mechanism of *coupled* dependence between spacer length and SC, and the quantitative contribution of the spacer length to the SC-sensitivity of the promoter can be computed without any adjustable parameter, as developed in the following.

The first prediction is the quantitative magnitude of this regulatory contribution. Geometrically, the size of the spacer (15-20 nt, in contrast to the shorter spacer of many dimeric TFs such as Fis or CRP, 8-10 nt) implies that physiologically relevant superhelical variations (of around *±*0.06) have an orientational effect on the - 10/-35 elements of the same magnitude as a variation in spacer length of one nucleotide (17 *×* 0.06 ≃ 1), and is therefore sufficient to induce a quantitative regulatory effect. In turn, this effect is tuned by the mechanical cost of aligning an ill-oriented spacer (e.g., the 18-nt spacer in the left panel of Fig. 1A), which depends on the experimentally known value of the torsional stifness of DNA [32]. Fig. 1B shows that, after computation, the addition/removal of one nucleotide in the spacer corresponds to a factor of around 2 in expression (e.g., compare the 18 vs 17 nt spacers at *σ* = *−*0.06). While this value is milder than the “on/off” regulation of many TFs affecting specific promoters, it is sufficient for a biologically relevant effect, especially if this factor affects RNAP activity in a global manner. This magnitude also approximately matches the observed effect of varying the spacer length (Supplementary Fig. S1). But interestingly, while the deformation energy has a symmetrical effect on the expression of spacers either too long or too short (same value of 16 and 18 nt spacers at *σ* = *−*0.06), the orientational effect of SC affects them in opposite directions, with short spacers being activated at relaxed levels (left) while long spacers are rather activated at highly negative levels (right).

### *In vitro* validation of model predictions on mutant promoters

In order to rigorously test the model, we developed an analysis of previously obtained *in vitro* transcription data involving spacer length mutants of model promoters. In this experimental protocol, plasmid templates containing *lacP* ^*s*^-derived promoters [45] and *PSC*-derived promoters [46] were prepared at well-defined superhelical densities and incubated with RNAP, thus minimising the effect of external regulatory factors. The raw datapoints (Fig. 2A) reflect not only the effect of the spacer, but also aforementioned regulatory contributions of SC, in particular the facilitated promoter opening explaining the overall (spacer length-independent) activation by negative SC [9]. We therefore normalised each datapoint by the one obtained at the same *σ* with the reference 17-nt spacer length (Fig. 2B), thereby eliminating these other factors and allowing a direct comparison with the model (by construction, the blue curve is flat; see the theoretical analysis in Materials and Methods, Eq. 5). The prediction is a linear dependence of the datapoints (coloured lines), where the *slopes* are set by the DNA torsional stifness value without any adjusted parameter, and proportional to (*n −* 17) (where *n* is the spacer length). In particular, the slope of the 19-nt spacer line is twice that of the 18-nt one (both activated by negative SC), and similarly for 15- and 16-nt spacers with negative values (repression by negative SC). Remarkably, the datapoints from two independent datasets obtained from different promoters (with 18- and 19-nt spacers) align along the predicted slopes without any adjustment, giving a strong support to our hypothesis that this regulatory contribution is dominated by the torsional elasticity of DNA (intercepts of the lines reflect promoter-specific features, see Materials and Methods).

In the study by Aoyama et al. [46], the same data were also collected with 16-nt spacers, but, in direct contrast to our predictions, these appear to be more activated at strong SC levels (Supplementary Fig. S5). Together with the previous ones, this observation suggests that our hypotheses are valid for some spacer lengths (18 and 19 nt), but fail for 16-nt spacers. Possible reasons and biological implications are developed in the Discussion, but briefly, the latter behaviour can be expected if their differential SC-sensitivity is dominated not by the deformation of DNA during closed-complex formation as we assume but, e.g., during open-complex formation. The question then arises, if this deviation is a feature of short spacers in general, thus strongly reducing the usefulness of our model, or if it is a mere exception of 16-nt spacers. We therefore ran additional *in vivo* experiments on mutant promoters containing opposite spacer lengths of 15 and 19 nucleotides.

### *In vivo* validation with opposite superhelical variations

We constructed spacer length mutants of the *pheP* promoter of *E. coli*, with a 17-nt (native), 15-nt (two deletions) or 19-nt spacer (two insertions, Supplementary Table S1). This promoter is SC-sensitive [47] and not regulated by any identified TF [48], and is thus a relevant candidate for this regulation mechanism based on the basal interaction with RNAP. Promoters were fused on plasmids in front of a luciferase reporter gene (Fig. 3A), and their expression was recorded in *E. coli* cells grown in LB rich medium in a microplate reader (Fig. 3B-C and Supplementary Fig. S3). Expression levels were computed shortly after treatment by the gyrase inhibitor novobiocin [3] or the topoisomerase I inhibitor seconeolitsine [55, 37] applied during exponential phase (see Materials and Methods). These drugs induce opposite SC variations in E. coli, DNA relaxation and overtwisting respectively [37], and thus provide complementary and independent tests of our predictions. Again, for a rigorous analysis of the data, we derived several parameter-free predictions (see dedicated paragraph in Materials and Methods, Eq. 7).

The native *pheP* promoter was repressed by novobiocin-induced DNA relaxation (Fig. 3B), in agreement with previous data [47], and was activated by seconeolitsine (Fig. 3C); both effects increased with the applied dosage. This result might be explained by the classical effect of SC-assisted promoter opening [9], as in the *in vitro* data above; the relatively modest (but highly significant) repression levels in response to novobiocin are due to a buffering effect of the reporter system (see Materials and Methods). Next, comparing the regulatory effect of the shocks on the mutants vs native promoter, we observed that novobiocin represses the 19-nt-spacer promoter with much stronger magnitude, whereas the 15-nt promoter was almost insensitive to DNA relaxation (Fig. 3F), following model predictions. Conversely, under seconeolitsine-induced DNA overtwisting, the activation fold-change was strongest for the 19-nt promoter, and very low for the 15-nt promoter (Fig. 3G). These four independent observations (differential effect of either superhelical variation on a shorter or longer spacer) were quantitatively reproduced by fitting the magnitude of the effective superhelical variations induced by the drugs (Fig. 3D-E, one fitted parameter for each shock, see Materials and Methods). These results confirm, after the *in vitro* data above, the prediction that 19-nt spacers are favoured by strong negative SC levels, and 15-nt spacers are rather favoured by DNA relaxation. The opposite behaviour of 16-nt spacers noted above thus seems to be an abnormal case where our hypotheses break down, and these promoters will therefore be disregarded in the upcoming analyses (see Discussion). For all other spacer lengths, since the proposed mechanism relies on the basal interaction between RNAP and promoter elements, we then wished to enlarge the analysis to the entire genome of *E. coli*, in order to test its validity and relevance at the global scale.

### Global effect of the spacer in promoter SC-sensitivity

We first looked at the variability of spacer lengths among *E. coli σ*70-dependent promoters, based on an available curated promoter map [48]. We focused on *σ*70 promoters, since it is predominant in exponential phase where the analysed samples were collected. The most frequent spacer length is 17-nt (27% total promoters) and most other are distributed between 15 and 19-nt (78% total promoters, Supplementary Fig. S6), and we therefore focused on that range in all analyses. Based on the results above, we hypothesised that promoters with short spacers would be more activated by DNA relaxation than those harbouring long ones (Supplementary Fig. S2). However, in contrast to the mutation studies above, genomewide analyses involve the comparison of promoters differing by many additional factors beyond their spacer, including genomic contexts, surrounding sequences, bindings of transcriptional regulators, etc. We thus searched for a statistical relation between promoter selectivity during SC variations and spacer length, rather than a prediction valid for all analysed promoters (see Materials and Methods).

In *E. coli*, the transcriptomic response to DNA relaxation was obtained with DNA microarrays [47], after a norfloxacin shock in two alternate topoisomerase mutant strains [56]. The analysis shows that promoters activated by DNA relaxation indeed harbour significantly shorter spacers than repressed ones (Fig. 4A, *P* = 0.007). Accordingly, classifying the promoters based on their spacer length (Fig. 4B) exhibits a clear decreasing tendency (correlation *P* = 0.07). The relatively high level of noise is due to the heterogeneity of promoter sequences within each group, and also likely to a fraction of inaccurately annotated promoters, since a single-nucleotide resolution in the definition of -10 and -35 hexamers is required for an accurate analysis but not always achieved, especially since the -35 element is not always well-defined [18]. As expected from the observations above, the 16-nt spacers again deviate from the model predictions (they were excluded from statistical analyses, see Discussion for a functional analysis of these promoters). For all others, these results suggest that, at the global scale, the variability of spacer length is used by bacterial cells for the selectivity of promoters in response to DNA relaxation.

### Analysis of transcriptomic data in various species

Since the investigated mechanism relies on highly conserved molecular actors, RNAP and topoisomerases, it might affect a particularly broad range of bacterial species, and we therefore wished to extend this analysis to other organisms. But while the transcriptomic response to DNA relaxation induced by gyrase inhibitors has been recorded in several species [3], curated and accurate promoter maps are generally lacking. We therefore based our analysis on available maps of transcription start sites (TSS) obtained from specifically designed transcriptomic data (Supplementary Tab. S2), followed by a scan for promoter motifs [54]. We thus obtained a list of putative promoters with associated *σ* factors and associated spacer lengths for two other enterobacteria, *Salmonella enterica* and the phytopathogen *Dickeya dadantii*, and at a drastically larger evolutionary distance, for the cyanobacterium *Synechococcus elongatus* and the small tenericute *Mycoplasma pneumoniae*. However, it must be noted that promoter prediction programs perform poorly in the detection of -35 elements: using the *E. coli* promoter map as a benchmark dataset, we found that the predicted -35 position deviated from the annotated one in around 50% promoters. In other species, this inaccuracy presumably resulted in a much higher level of statistical noise than for the annotated *E. coli* promoters above. In spite of this difficulty, a difference of spacer length in the same direction as in *E. coli* was observed in all investigated species (Supplementary Fig. S4), albeit with weaker magnitudes and levels of statistical significance. Altogether, while improvements in promoter definition are clearly required for a solid conclusion, this systematic observation suggests that the variability of spacer length might indeed underpin a selective activation and repression of promoters by global SC variations throughout the prokaryotic kingdom.

### Inheritable selection of promoters based on the spacer length

We finally investigated if the present mechanism could be involved not only in transient DNA relaxation responses induced by antiobiotic shocks, but also in inheritable variations of global gene expression in the longest-running evolution experiment [57, 58]. Indeed, in this experiment involving the growth of *E. coli* cells in a daily refreshed minimal medium, point mutations affecting the SC level were quickly and naturally selected as they provided substantial fitness gains [57]. A first mutation (in *topA*) was fixed before 2,000 generations, and a second mutation (in *fis*) before 20,000 generations, both leading to an inheritable increase of negative SC (Fig. 5A). Based on our modelling and the previous observations, we therefore expected promoters with a long spacer to experience enhanced expression in the evolved strains compared to the ancestor. Such a relation is indeed observed, both after 2,000 generations (Fig. 5B, *P* = 0.04) and 20,000 generations (Fig. 5C, *P* = 0.026). The signal is significant but slightly weaker than that observed with antibiotics (Fig. 4); this may be explained by the inheritable (rather than transient) nature of the SC variation, which induces an adaptive response of the cells via other regulatory pathways. Again, these results suggest that promoters of different spacer lengths respond differently to SC variations due to, e.g., mutations in topoisomerase genes that are observed even between closely related species [59].

## Discussion

While the spacer length and sequence are known to modulate transcriptional activity, we wished to quantitatively model and test the long-proposed idea [45, 46, 10] that SC plays a specific role in this process through a simple and basal orientational effect during closed-complex formation. This effect indeed emerged as a predictable quantitative signal, both in specifically designed mutant promoter assays, and as a statistical tendency in whole-genome data. The model and the latter results altogether suggest that this mechanism has a widespread relevance in bacterial transcription, although more detailed and comprehensive analyses will be required to confirm it in various species.

### Limitations of the regulatory model: the case of 16-nt spacers

The model was based on the hypothesis that the investigated mechanism (torsional orientation of -10 and -35 binding sites) could be decoupled both from further regulatory effects of SC (assumed to be independent of the spacer length), and from further modulating effects of the spacer length and sequence (assumed to be independent of SC). These hypotheses, already proposed and supported by a collection of qualitative observations on individual promoters [10], are validated here *a posteriori* by the quantitative agreement between model predictions and analysed data of different kinds. A notable exception, however, are promoters involving 16-nt long spacers, for which all observations converge to an opposite behaviour resembling that of long spacers (Supplementary Fig. S5, Fig. 4B). This behaviour does not imply that the mechanism does not occur for this family of promoters, but possibly that its effects are overcome by a stronger opposite regulatory effect of SC at a later stage of transcription, in particular during open-complex formation where different elements of the promoter make extensive and complex contacts with RNAP, and the destabilisation of the double helix by SC has a drastic influence on the expression level [60, 16, 9].

Interestingly, a well-studied class of promoters involving 16-nt spacers are those encoding stable RNAs in *E. coli*, subject to stringent control. These experience a strong repression by ppGpp as well as DNA relaxation occurring in the cell upon transition to stationary phase [61, 62, 63]. Both effects were attributed to the unusual kinetics of promoter opening and escape due to their G/C-rich “discriminator” sequence inducing unusual interactions with RNAP in the open-complex [64, 60, 65]. We note however that, based on the observations above, the repressive effect of DNA relaxation is also favoured by the 16-nt long spacer of these promoters (Supplementary Fig. S5, Fig. 4), and both elements (discriminator and spacer) might thus contribute to their behaviour. To test this hypothesis further, we looked if a similar relation exists among promoters of protein-encoding genes. Indeed, we found that those containing 16-nt spacers exhibit significantly G/C-richer discriminators compared to all other promoters (Supplementary Fig. S7, *P <* 0.001), suggesting a tight relation between both properties (spacer length and discriminator sequence). The subclass of 16-nt long spacer, G/C-rich discriminators might thus experience a specific pathway in open-complex formation, beyond the range of our model focused on the closed-complex. This notion is further emphasised by the observation that the reactivity of spacer DNA to potassium permanganate or DMS within the open-complex depends on the spacer length or sequence, indeed suggesting an effect at that later stage of transcription [28, 66].

### Additional factors influencing the relative orientation of -10/-35 elements

While we focused on the effect of SC on the direct interaction of RNAP with promoter DNA, the (un)twisting of the spacer has been proposed as the mechanistic basis for the regulatory action of several TFs, including MerR, which regulates the (*mer*) operon encoding components of the mercury (Hg) resistance system [31]. The *mer* promoter includes a MerR binding site overlapping -35 and -10 elements, which are separated by a 19-nt spacer, and this unusual length was shown to be essential for normal activation by MerR [67]. In the absence of Hg(II), MerR binds to the *mer* promoter in its repressor conformation. The presence of Hg(II) causes a conformational change in MerR and, in turn, the untwisting of DNA and reorientation of -35/-10 sites for effective open-complex formation [31]. In agreement with this model, a global increase in negative SC facilitates MerR-mediated activation, impedes MerR-mediated repression, and conversely for DNA relaxation [68]. The same mechanism presumably applies to other metal-dependent regulators[31, 69], and to the activation of the 19-nt spacer *mom* promoter by the C protein of bacteriophage Mu [70]. The general elastic model proposed here may thus be enriched to include these more specific actors.

### Effect of spacer sequence on -35/-10 alignment by RNA Polymerase

Apart from the limitations mentioned above, we only considered how SC modulates the relative torsional orientation of -10 and -35 elements depending on the spacer length, and neglected any effect of its base sequence. However, the latter has been shown to affect transcriptional activity of various promoters [26, 22, 23, 71, 24, 72, 25]. Focusing specifically on the torsional orientation between the -35 and -10 sites (i.e., the geometrical parameter most sensitive to SC), the spacer sequence might modulate two parameters that were considered as constant in our equations: the average twist angle between successive nucleotides (*α*_0_ in Eq. 1) and the torsional stifness (*k*_*θ*_), whose values were estimated for all basepair step sequences from a collection of crystallographic structures [32]. We implemented these values [33] to estimate the magnitude of the resulting adjustment of the regulatory effect of SC, for all *σ*70-dependent promoters of *E. coli* (Supplementary Fig. S8, with details of the computation in Supplementary Information). Over-all, the maximal span of the sequence contribution remains weaker than the gain/loss of one nucleotide in the spacer (Supplementary Fig. S8A), confirming our hypothesis that the effect of the spacer sequence is weaker than that of its length for the considered orientational angle. More precisely, we then wondered if this modulation results from the sequence-induced heterogeneity of the twist angle (i.e., the DNA structure) or stifness (i.e., elasticity), or both. We therefore modified the computation to impose a sequence-averaged value to either of these two parameters (Supplementary Figs. S8 B-C). We observed that the effect of the sequence on DNA stifness alone has almost no regulatory effect (Supplementary Fig. S8B), meaning that the sequence contributes mostly by modulating the total twist angle of the spacer, i.e., its average structure rather than its elasticity. As noted, this modulation yet remains modest compared to that induced by the variability in spacer length.

## Acknowledgements

We thank the whole CRP team, Ralf Everaers and Ivan Junier for helpful discussions, and Nicolas Paulhan for experimental contributions.

