## Supplementary Information for "Quantitative contribution of the spacer length in the supercoiling-sensitivity of bacterial promoters"

February 24, 2022

**This PDF file includes:**

- Supplementary text
- Supplementary figures S1 to S8
- Supplementary tables S1 to S2
- Supplementary information references

### Supplementary text

#### Genome-wide analyses of spacer responses to supercoiling variations in distant bacteria

Transcriptomic data were collected from the literature (Supplementary Tab. S2). Since curated promoter maps were not available, genome-wide TSS maps and associated genes were retrieved from literature (Supplementary Tab. S2), and a scan for promoter motifs upstream of TSSs was conducted with bTSSfinder [1]. Only $\sigma$70-dependent promoters were retained for all organisms, except for the cyanobacterium *S. elongatus*, for which only $\sigma$A-dependent promoters were kept (primary $\sigma$ factor), and classified depending on their spacer length and response to the investigated condition. The thresholds for statistical selection procedures are indicated in Supplementary Tab. S2, and were adjusted to generate subsets of act/rep genes of sizes comparable among the different data sets, while having enough statistical power for the analysis. The relation between promoter activation and spacer length was then quantified by a Student's t-test between activated and repressed promoters (Supplementary Fig. S4), such as in Fig. 4 and 5. All error bars shown are 95% confidence intervals.

#### Modelling of the effect of spacer sequence on -35/-10 alignment by RNA Polymerase

To compute the contribution of the spacer sequence in the model (Discussion), we included the sequence-dependence of two parameters of Eq. 1, the twist angle ($\alpha_0$) and the torsional stiffness ($k_\theta$), based on the parameters of the rigid base-pair model inferred from high-resolution crystallographic structures of DNA oligomers [2] and implemented in TwistDNA [3]. Based on these parameters, Eq. 1 was modified to include the total angle between -35/-10 binding sites and the total spacer stiffness, resulting in the following:

$$\Delta G(\sigma, s) = \frac{1}{2} k_\theta(s) \left( \theta_P - \theta_0(s) - \alpha_0 \frac{k_\theta}{k_\theta(s)} \sigma \right)^2$$

where $\sigma$ is the supercoiling level, $s$ is the promoter spacer sequence, $k_\theta(s)$ is the DNA spacer twist stiffness estimated with ThreaDNA [3], $\theta_P = 543$ is the optimal angle between -35 and -10 sites for RNAP binding, as assumed in the main text (with most predictions given in Discussion being independent of its value), $\theta_0(s)$ is the effective angle estimated with ThreaDNA [3] which depends on base composition, $\alpha_0 = 34°$ is the average twist angle between adjacent nucleotides, and $k_\theta$ is the average (base-pair step) twist stiffness. Then, for each *E. coli* $\sigma$70-dependent promoter, the response to DNA relaxation was predicted starting from a level $\sigma = -0.06$, with a relaxation magnitude $\Delta\sigma = 0.03$ (Supplementary Fig. S8).

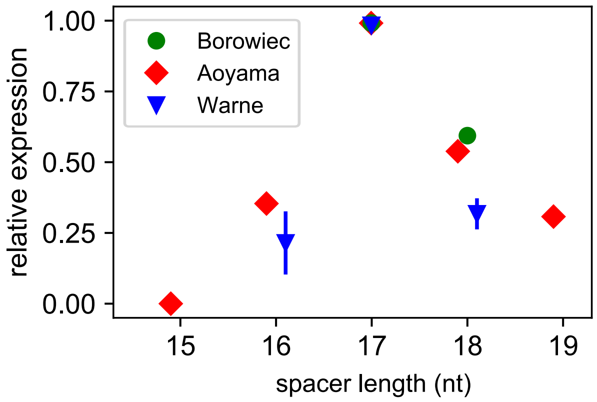


**Supplementary Figure S1**: Relative *in vitro* promoter expression levels depending on spacer length, for a supercoiling level $\sigma = -0.06$, using data from [4] (green), [5] (red), and [6] (blue). Error bars are shown for [6] data due to the usage of different promoters for which no SC level was measured.

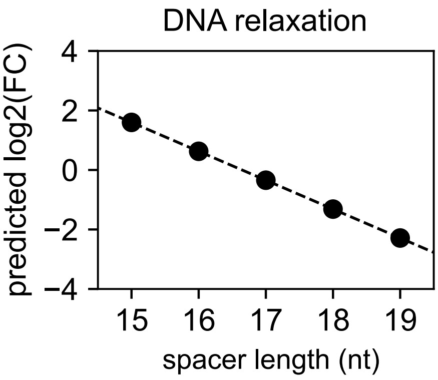


**Supplementary Figure S2**: Predicted expression fold-changes for promoters containing 15 to 19-nt spacers, during a relaxation from $\sigma = -0.06$ to $\sigma = -0.03$.

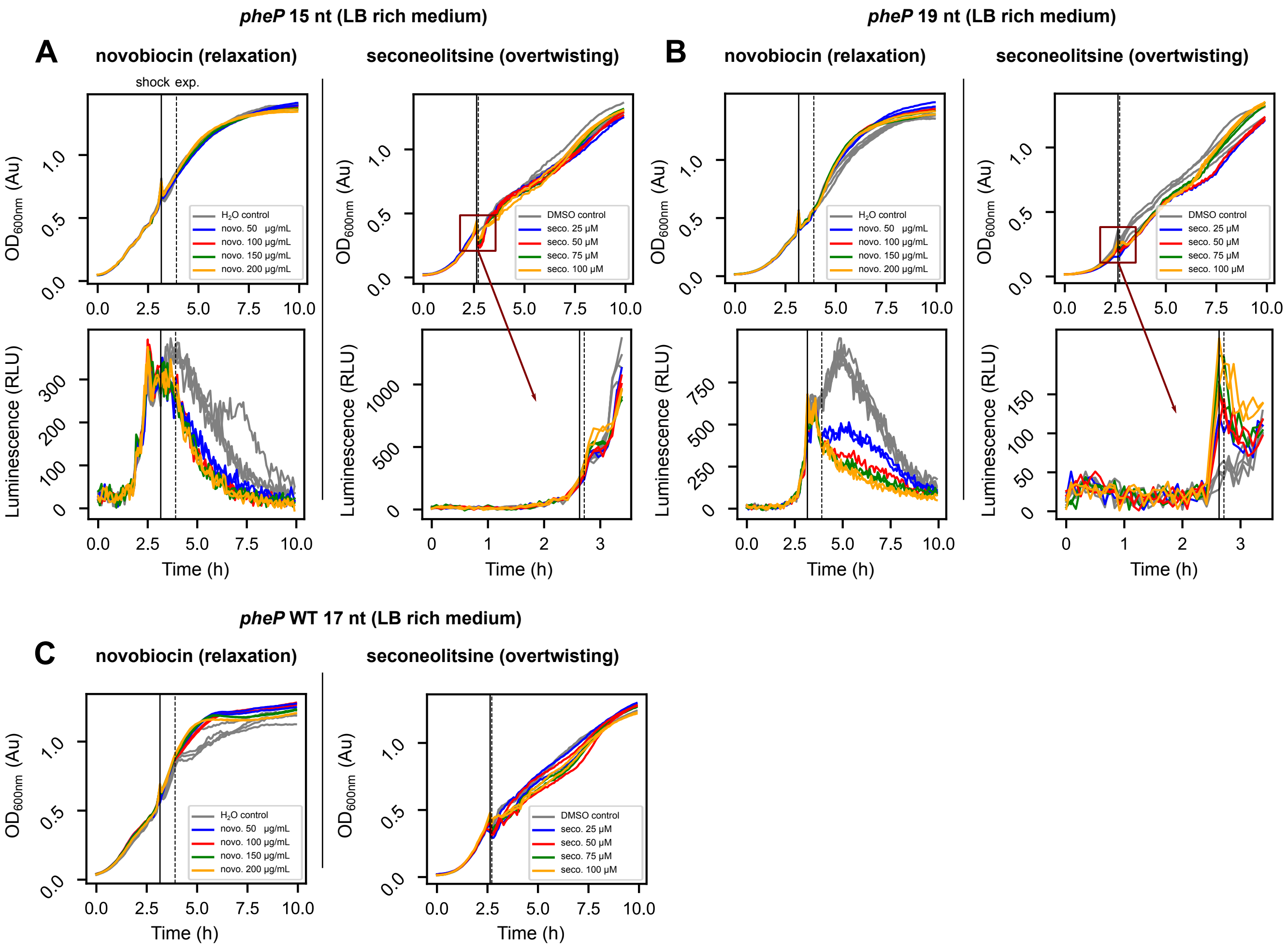


**Supplementary Figure S3**: Bacterial growth and promoter expression monitored in a microplate reader (raw data). **(A)** *E. coli* bacteria carrying plasmids with the *pheP* mutant promoter with a 15-nt spacer. **(B)** *pheP* mutant promoter with a 19-nt spacer. **(C)** *pheP* native promoter containing a 17-nt spacer. For all promoters, upper panels show the bacterial growth measured by $OD_{600nm}$, lower panels the expression levels measured by luminescence, left panels the experiments conducted with novobiocin, and right panels the experiments conducted with seconeolitsine. "Shock" and "exp." refer to the time points used for the antibiotic shock, and for the computation of expression fold-changes, respectively. Due to the stronger and more transient expression variation induced by seconeolitsine shock, a zoom is performed for luminescence. Some $OD_{600nm}$ curves exhibit slight discrepancies at shock time or after several hours; these are optical artefacts due to the opening of the recorder and/or the formation of sediments disrupting the measurements. The luminescence curves of *pheP* native 17-nt spacer promoter are provided in Fig. 3.

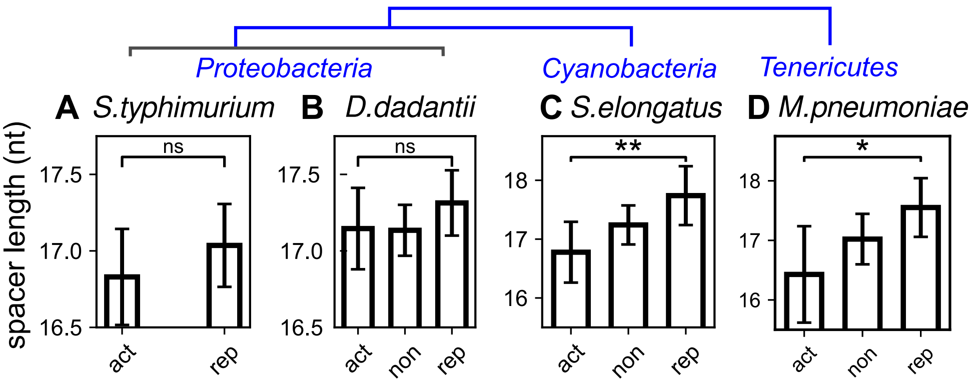


**Supplementary Figure S4**: Statistical relation between spacer length and promoter's response to DNA relaxation (act: activated, non: not significantly affected, rep: repressed) in distant bacterial species (see Supplementary Tab. S2 for experimental conditions and references): **(A)** *Salmonella typhimurium* ($P = 0.169$), **(B)** *Dickeya dadantii* ($P = 0.170$), **(C)** *Synechococcus elongatus* ($P = 0.007$), **(D)** *Mycoplasma pneumoniae* ($P = 0.017$). In proteobacteria and *M. pneumoniae*, only $\sigma 70$ promoters were considered, whereas only $\sigma A$ promoters were considered for *S. elongatus*. A schematic phylogeny is depicted above.

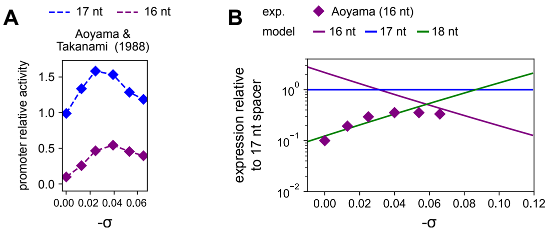


**Supplementary Figure S5**: Comparison of *in vitro* transcription assays for 16-nt spacers to model predictions. **(A)** Relative activities of promoters with 17 and 16-nt spacers (blue and purple, respectively) depending on supercoiling level $\sigma$, data from [5]. **(B)** Transcription model predictions (solid lines) compared to the *in vitro* promoter expression data (exp.). Each datapoint was normalised by the corresponding value obtained in the reference mutant promoter with a 17-nt spacer (see text). The (weakly) positive slope of the datapoints corresponds to the behaviour expected for long rather than short spacers (green line for the prediction corresponding to a 18-nt spacer).

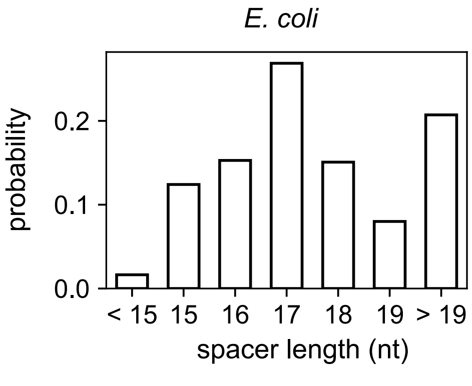


**Supplementary Figure S6**: Distribution of spacer lengths at *E. coli* $\sigma 70$-dependent promoters, based on the Ecocyc database [7].

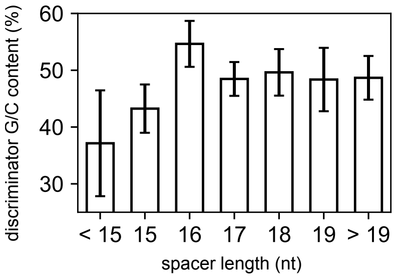


**Supplementary Figure S7**: Analysis of the coupling between promoter spacer length and discriminator G/C-content, based on the Ecocyc database [7]. Promoters with 16-nt spacers exhibit G/C-richer discriminators compared to other promoters ($P$-value < 0.001), 15-nt and < 15-nt spacers exhibit A/T-richer discriminators ($P$-value = 0.015 and 0.004, respectively), whereas 17, 18, 19 and > 19-nt spacers have similar, not significantly different discriminator A/T-contents.

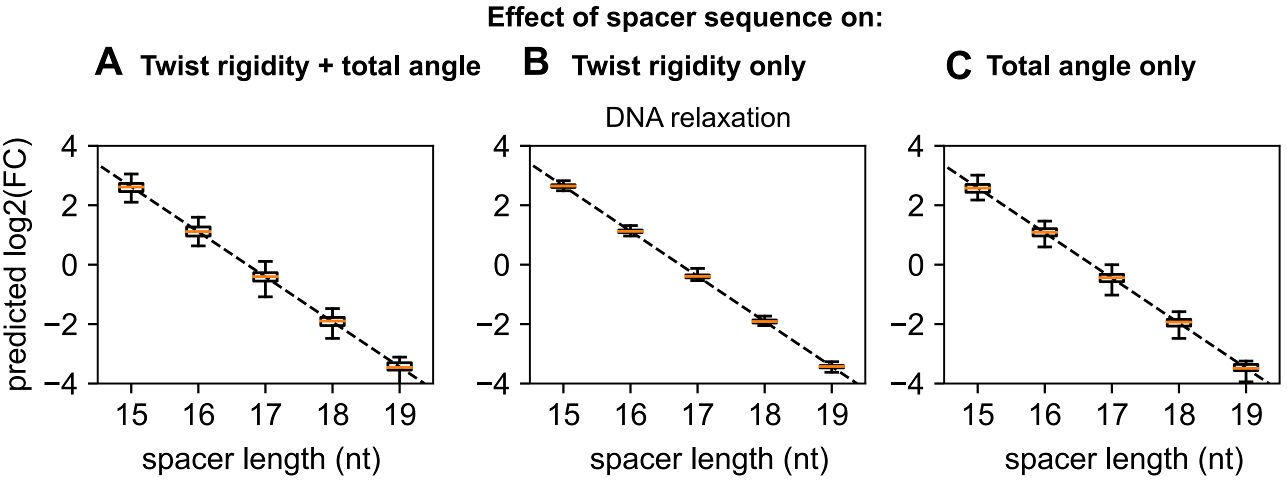


**Supplementary Figure S8**: Effect of the spacer sequences: boxplots show the span of predicted fold-changes for all $\sigma$70-dependent promoters of *E. coli* (based on the Ecocyc database [7]), during a relaxation shock from $\sigma = -0.06$ to $\sigma = -0.03$. The box extends from the first quartile to the third quartile values of the data, with an orange line at the median, and with whiskers extending from each end of the box to the extreme values. **(A)** With sequence-dependent values for both twist angle and rigidity: the maximal magnitude of the sequence effect is comparable to that of a gain or loss of one nucleotide in the spacer. **(B)** With sequence-dependent spacer twist rigidity and sequence-averaged total angle, the effect of the sequence is strongly reduced, showing that the latter mostly modulates the angle. **(C)** With sequence-dependent total angle and sequence-averaged spacer twist rigidity.

| **Promoter** | **Sequence** |
| --- | --- |
| *>pheP_WT_17nt* | CTCGAGTCAGAGGTGATGAGCCGGATTGCCGCGCCGATGATTGGCGGCATGATCACCGCACCTTTGCTGTCGCTGTTTATT<br>ATCCCGGCGGCGTATAAGCTGATGTGGCTGCACCGACATCGGGTACGGAAATAAAAGCAGGATACCCCGTTTAACCGTGTG<br>GATTGTGTCTTGCGA**CGATGGGCACTAAATGT**TAAAAGGTGCCCCTCAACAAAAAAGACACACAGGGGAAAGGCGGATCC |
| *>pheP_15nt* | CTCGAGTCAGAGGTGATGAGCCGGATTGCCGCGCCGATGATTGGCGGCATGATCACCGCACCTTTGCTGTCGCTGTTTATT<br>ATCCCGGCGGCGTATAAGCTGATGTGGCTGCACCGACATCGGGTACGGAAATAAAAGCAGGATACCCCGTTTAACCGTGTG<br>GATTGTGTCTTGCGA**CGATGGGCTAAATGT**TAAAAGGTGCCCCTCAACAAAAAAGACACACAGGGGAAAGGCGGATCC |
| *>pheP_19nt* | CTCGAGTCAGAGGTGATGAGCCGGATTGCCGCGCCGATGATTGGCGGCATGATCACCGCACCTTTGCTGTCGCTGTTTATT<br>ATCCCGGCGGCGTATAAGCTGATGTGGCTGCACCGACATCGGGTACGGAAATAAAAGCAGGATACCCCGTTTAACCGTGTG<br>GATTGTGTCTTGCGA**CGATACGGGCACTAAATGT**TAAAAGGTGCCCCTCAACAAAAAAGACACACAGGGGAAAGGCGGATCC |
| | XhoI restriction site, -35 element, **spacer**, -10 element, TSS, BglII restriction site |

| **Plasmid** | **Description** | **Origin** |
| --- | --- | --- |
| pUCTer-*luc* | High-copy-number vector (pUC18 derivative) containing a multiple cloning site upstream of the *luc* reporter gene, followed by a *rrnB* terminator and a *cat* gene conferring chloramphenicol resistance. | Laboratory collection |

**Supplementary Table S1**: List of *pheP*-derived synthetic promoter sequences with mutated spacers, and pUCTer-*luc* plasmid used in this study.

| **Species** | **Condition** | **Threshold adj. P-value** | **Threshold $\log_2$(FC)** | **Supercoiling variation** | **Transcription start sites reference** |
| --- | --- | --- | --- | --- | --- |
| *Escherichia coli* | norfloxacin [8] | 0.05 | 0 | relaxation | [7] |
| | inheritable supercoiling variation [9] | 0.1 | 0 | overtwisting | [7] |
| *Salmonella typhimurium* | *gyrA* mutant [10] | NA | 0.4 | relaxation | [11] |
| *Dickeya dadantii* | novobiocin [12] | 0.05 | 0 | relaxation | [13] |
| *Synechococcus elongatus* | supercoiling correlation* [14] | NA | 0.4 | relaxation/ overtwisting | [15] |
| *Mycoplasma pneumoniae* | novobiocin [16] | 0.1 | 0 | relaxation | [16] |

**Supplementary Table S2**: Compilation of investigated species, conditions and references. The thresholds for statistical selection procedures are indicated, and were adjusted to generate subsets of activated/repressed genes of sizes comparable among the different data sets. For a threshold of 0.4 on the $\log_2$(fold-change), genes are considered activated for a $\log_2(FC) > 0.4$, repressed for a $\log_2(FC) < -0.4$, and not significantly affected for a $\log_2$(FC) comprised between -0.4 and 0.4. The correlation* condition from *S. elongatus* corresponds to the phasing of gene expression in the SC circadian oscillation and provides an indirect proxy of gene response to SC relaxation [14].
